# Effect of spatial heterogeneities on minimal stochastic models of cell polarity

**DOI:** 10.64898/2026.03.27.714696

**Authors:** Valentin Anfray, Hong-Yan Shih

## Abstract

Asymmetric self-organization is a hallmark of cell polarity, yet the diversity of observed polarization patterns is frequently attributed to specialized, complex biochemical mechanisms motifs beyond simple positive feedback. Here, we demonstrate that spatial heterogeneity alone fundamentally reshapes polarization dynamics within minimal stochastic reaction–diffusion processes. We show that weak differences in reaction rates between distinct spatial domains strongly bias polarization timing and determine which region ultimately polarizes. In systems containing two distant favored regions, a “stochastic winner-takes-all” mechanism—driven by long-range competition mediated by a shared cytoplasmic pool—induces stochastic switching that manifests as pole-to-pole oscillations. By relaxing the assumption of a perfectly mixed cytoplasm and incorporating finite cytoplasmic diffusion, we reveal a qualitative shift in this competitive dynamic. Specifically, as the total particle abundance increases, the system transitions from monopolar to bipolar activation, capturing the essence of the New-End Take-Off (NETO) phenomenon during cell growth and provides a physical basis for pole coexistence. These results demonstrate that spatial heterogeneity alone can strongly influence polarization dynamics in minimal models, highlighting the potential importance of quenched spatial variability in biological reaction–diffusion systems.

**Author summary:** Cells often need to choose a specific site for growth, division, or shape change. This process, known as cell polarization, is a fundamental organizing principle in biology. The wide variety of polarization patterns seen in living cells is often explained by proposing complex biochemical mechanisms beyond basic positive feedback among signaling molecules. In this work, we asked whether some of this diversity could instead arise from a simpler source: fixed spatial differences within the cell. Using minimal stochastic reaction-diffusion models, we found that even small local differences can strongly influence where polarization appears and how quickly it develops. When two favored sites are present, they can compete for a shared pool of molecules in cytoplasm, so that one site dominates at a time and the polarized state can switch stochastically between them. We also found that this competition changes when the shared molecular pool does not mix instantly: under these conditions, two polarized sites can start to coexist. This behavior offers a simple physical explanation for phenomena such as the appearance of a new growth site during cell development. Our results show that spatial heterogeneity alone can generate behaviors that might otherwise seem to require much more complicated biochemical mechanisms.

## Introduction

Cell polarity is a fundamental organizing principle of living systems, underlying processes such as neuronal morphogenesis [1, 2], cell migration [3], directional cell growth [4], asymmetric cell division [5], and tissue development [6, 7]. Polarity refers to the emergence and maintenance of spatial asymmetry in cell morphology and intracellular protein distributions [8]. In many organisms, polarization is controlled by Rho-family GTPases (such as Cdc42, Rac, or Rop) [9–11]. A widely accepted, but no unique, mechanism for sustaining this polarization is the existence of a positive feedback loop that promotes the local GTPase activation [12, 13].

To understand the minimal requirements for spatial pattern formation—such as the emergence of polarized domains—reaction–diffusion models, building on the pioneering work of Turing [14] and Gierer and Meinhardt [15], have been extensively utilized [16]. In the context of cell polarity, both deterministic and stochastic reaction–diffusion models have been shown to reproduce polarization dynamics and robustness properties [17–19]. In particular, mass-conserved models [17, 20–24] have proven highly effective for modeling cell polarity. Rather than producing the multi-domain patterns characteristic of classical Turing instabilities, these models naturally account for the emergence of a single polarity axis or a limited number of poles. In such systems, the strong disparity in diffusion rates between slowly diffusing membrane-bound (active) proteins and their rapidly diffusing cytosolic (inactive) counterparts drives a mechanism of local activation coupled to global depletion.

Although spontaneous polarization can arise from intrinsic nonlinear dynamics, these studies generally assume spatially uniform rates and environment, despite the fact real cells are rarely perfectly homogeneous. Pre-existing spatial cues—such as external chemoattractant gradients [25], membrane curvature effects [26–29], or cortical landmarks involved in bud-site selection [10, 30–32]—explicitly break translational symmetry and bias polarity establishment. By contrast, the possible role of intrinsic spatial heterogeneity, or quenched disorder, has been comparatively little explored. In other fields, quenched disorder is known to qualitatively alter macroscopic behavior such as in Anderson localization [33], spin glasses [34] and ecological dynamics [35, 36]. In the realm of critical phenomena, scaling arguments like the Harris [37] and Imry–Ma [38] criteria successfully predict whether uncorrelated spatial inhomogeneities will alter the critical properties of a pure system. Even away from the critical point, large rare disorder fluctuation can cause nonanalyticities, known as Griffiths singularities [39, 40]. Although biological environments exhibit more rapid temporal fluctuations compared to in solid state physics, these fluctuations can effectively act as frozen (quenched) heterogeneities when evaluated against the characteristic timescales of processes such as budding or cell division [41, 42].

Interestingly, spatial heterogeneity is already implicitly present in several models of cell polarity, particularly in studies of oscillatory dynamics in rod-shaped cells. In these systems, the cell is often represented as two spatially separated compartments—typically corresponding to the two tips of the cell—coupled through cytoplasmic diffusion [43–48]. Other approaches introduce position-dependent reaction rates, for instance by allowing activation rates to decay exponentially with the arc-length distance from the nearest cell tip [49]. Such compartmental descriptions successfully capture oscillatory localization of polarity proteins observed in organisms such as fission yeast [11, 50, 51]. More generally, oscillatory or switching polarity dynamics can arise when additional ingredients are incorporated, including delayed feedback [50], negative feedback [52], or biased transport processes [53]. A related phenomenon is the New-End Take-Off (NETO) transition observed in certain rod-shaped cells [50, 54], where cell growth triggers a switch from monopolar to bipolar growth. This transition is often attributed to saturation mechanisms that stabilize two polarized sites once the total number of polarity proteins becomes sufficiently large [44, 50]. Similar mechanisms have also been invoked to explain the coexistence of multiple polarization sites in more general contexts [55, 56]. These studies illustrate how spatial asymmetry and compartmentalization can qualitatively influence polarity dynamics. At the same time, they raise a broader question: while stochastic models can generate spontaneous cell polarization with minimal ingredients [57], whereas deterministic descriptions typically require additional regulatory mechanisms [18, 19], it remains unclear what ingredients are necessary for the emergence of oscillatory dynamics or NETO-like transitions.

Recent studies of the diffusive epidemic process (DEP)—a minimal stochastic reaction–diffusion system characterized by two diffusing species and positive feedback—have demonstrated that quenched spatial disorder can dramatically alter its critical behavior [58]. Specifically, the presence of disorder leads to significantly slower dynamics, driven by the emergence of local regions that remain polarized for exceptionally long durations. Because the DEP shares structural similarities with minimal models of spontaneous polarization [57], these findings suggest that spatial heterogeneity could qualitatively modify the dynamics of polarity establishment.

Motivated by these considerations, we investigate how quenched spatial heterogeneities affect a minimal stochastic reaction–diffusion model capable of spontaneous polarization, as well as the DEP away from its critical regime. We introduce both processes and derive simplified variants that facilitate analytical study. We first analyze polarization dynamics in a system composed of two spatial regions and quantify how polarization times of each region depend on the total particle number and their relative favorability. Then, we extend this analysis to two spatially separated but equally favored regions and show that quenched heterogeneity can induce stochastic switching between competing polarization sites. We relax the assumption of a perfectly homogeneous cytoplasm and demonstrate that multiple polarized zones may stably coexist. Finally, we incorporate uniaxial cell growth and show that growth can drive a transition from a single, potentially switching, polarized pole at one tip to a regime where stable polarization at both tips emerges.

## Models

### Neutral Drift Polarity (NDP) Model

We first consider the neutral drift polarity (NDP) model [57, 59], which was introduced to explain the spontaneous emergence of cell polarity from minimal ingredients; see Fig. 1a). In this model, cytosolic particles, denoted *c*, are assumed to be homogeneously distributed. This assumption is justified by the large separation of proteins diffusivities between the cytoplasm and the membrane [16, 60]. Cytosolic particles can spontaneously bind to the membrane at rate *k*_*on*_. Once bound, membrane-bound particles, denoted *m*, diffuse along the membrane with diffusion coefficient *D*, can unbind at rate *τ*, and can recruit additional cytosolic particles at rate *λρ*_*c*_, where *ρ*_*c*_ denotes the local density of cytosolic particles sensed by a membrane-bound particle. A key property of this model is the spontaneous formation of a polarized cluster, defined as a majority of particles *m* localized within a finite fraction of the system, when the total particle number exceeds a critical value. Clustering is lost again when the total particle number becomes too large [59].

**Fig 1.**
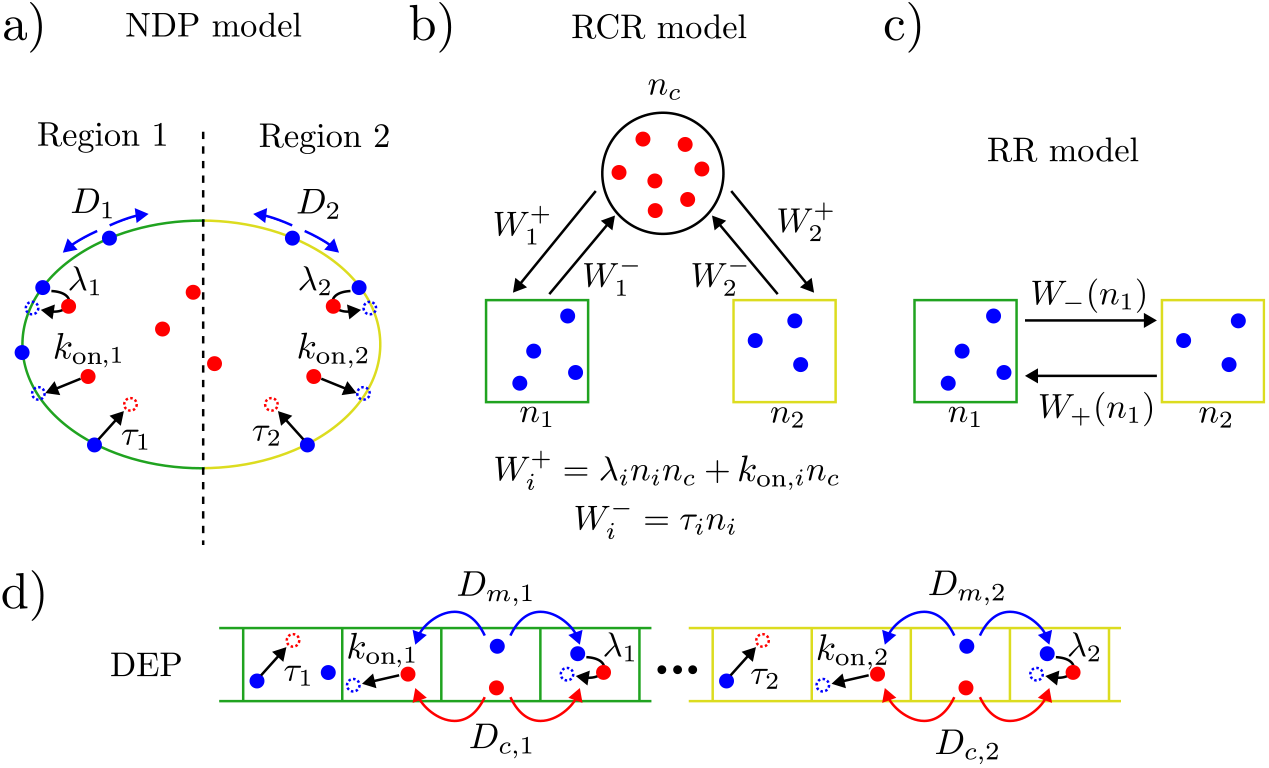
Schematic representation of the reaction-diffusion models considered in the presence of quenched spatial disorder. Two spatial regions (green and yellow) are characterized by distinct local parameters *D*_*i*_, *λ*_*i*_, *τ*_*i*_, and *k*_on,*i*_. Blue circles represent membrane-bound particles, while red circles denote particles in the cytosol. (a) In the NDP model, membrane-bound particles diffuse along the membrane with a spatially dependent diffusion coefficient *D*_*i*_, while the cytosolic density remains homogeneous. (b) The RCR model simplifies the NDP by neglecting membrane diffusion, focusing instead on the fluctuations of particle numbers within each region and the cytosol. (c) The RR model introduces a further simplification by conserving the total number of membrane-bound particles. (d) The DEP model is a discrete formulation that explicitly accounts for particle diffusion within the cytosol in an simple way.

Our main interest lies in understanding the effect of spatially quenched disorder. To this end, we consider a system composed of *k* spatial regions characterized by distinct local parameters *τ*_*i*_, *λ*_*i*_, *D*_*i*_, and *k*_*on,i*_ with *i* ∈ [1, *k*]. Numerical simulation details are provided in Appendix S3 Numerical simulations.

### Region-Cytoplasm-Region (RCR) Model

Assuming that these regions are sufficiently large—or equivalently that membrane diffusion is sufficiently slow—a first approximation is to treat them as spatially disconnected subsystems that interact solely through exchanges with the cytoplasm. Furthermore, we neglect explicit spatial degrees of freedom within each region, since membrane diffusion does not affect the total number of particles in a given region. This leads to a reduced description, hereafter referred to as the Region–Cytoplasm–Region_*k*−1_ (RCR_*k*−1_) model. An illustration of this framework for *k* = 2 (denoted simply as RCR in the following) is provided in Fig. 1b). We denote by *n*_*i*_ the number of membrane-bound particles in region *i*, and by *n*_*c*_ the number of cytosolic particles. Particle conservation imposes

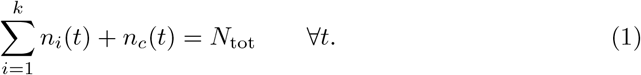

The local parameters in region *i* are denoted by *λ*_*i*_, *τ*_*i*_, and *k*_*on,i*_. The stochastic dynamics is governed by the following microscopic transitions:

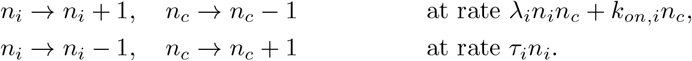

The corresponding deterministic mean-field equations read

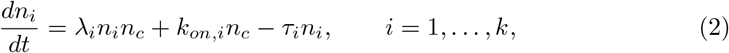

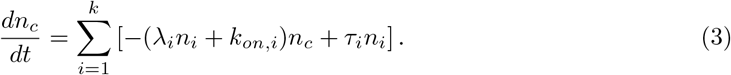

At stationarity, the membrane populations satisfy

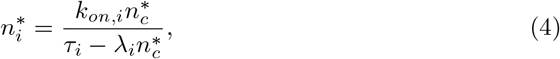

where 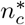 is obtained from the conservation law 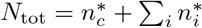. This yields a polynomial equation of order *k* + 1 for 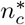. A unique physical solution is selected by the constraints 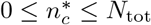 and 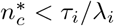 for all *i*, ensuring 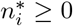.

### Region-Region (RR) Model

To obtain an explicit expression for the stationary distribution, we further simplify the problem by assuming that the total number of membrane-bound particles,

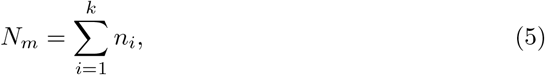

is constant. We focus on the case *k* = 2, which is the only configuration for which a closed-form stationary distribution can easily be derived. In this reduced description, a particle unbinds from region 1 at rate *τ*_1_*n*_1_ and from region 2 at rate *τ*_2_(*N*_*m*_ − *n*_1_).

Upon unbinding, the particle immediately rebinds to region 1 with probability

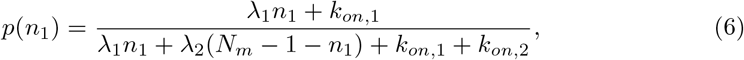

and to region 2 with probability 1 − *p*(*n*_1_).

The transition rates for increasing or decreasing *n*_1_ by one particle are therefore given by

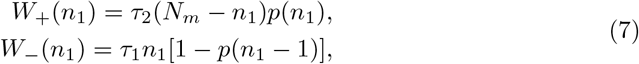

with the boundary conditions *W*_−_(0) = 0 and *W*_+_(*N*_*m*_) = 0 by construction.

The resulting Region–Region (RR) model, illustrated in Fig. 1c), is described by the master equation

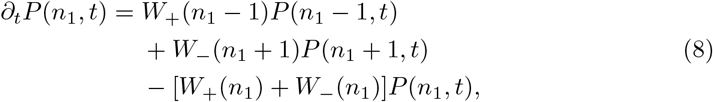

where *P* (*n*_1_, *t*) is the probability that region 1 has *n*_1_ particles at time *t*. As the number of particle can only increase or decrease by one, such system corresponds to a birth-and-death process for which many analytical results are known. In particular, birth-and-death processes admit a stationary distribution *P*_*s*_ of the form [61]

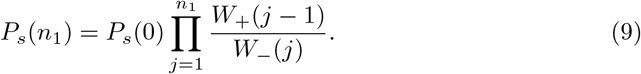

### Diffusive Epidemic Process (DEP)

A central assumption of the NDP model, and hence of the RCR and RR models, is the homogeneity of the cytosolic particle distribution. As we will show later, this assumption breaks down when the local density of membrane-bound particles becomes large, i.e., when the reaction timescale in a given region exceeds the typical time the cytosolic particles need to diffuse in the system. To relax this assumption, we discretize space and explicitly model the diffusion of cytosolic particles, treating them as an additional diffusing species on a lattice. This framework corresponds to the diffusive epidemic process (DEP) [62, 63], as illustrated in Fig. 1d). Additional details are provided in Appendix S2 DEP, where we discuss its relation to the NDP model, and in Appendix S3 Numerical simulations, where we describe the numerical simulation procedure.

In the following, except status otherwise, values of the parameters used in the numerical simulation are listed in Appendix S4 Parameters.

### Effect of disorder on two regions

In the context of cell polarity, quenched disorder, manifesting as spatially localized heterogeneities, is to be expected in many scenarios, such as in the presence of a pheromone concentration gradient in mating fission yeast cells [64], or more generally due to chemical gradients [25], or variations in membrane curvature [26–29] or bud-site-selection landmarks [10, 30–32]. While determining the microscopic effect of these heterogeneities is a complex biological problem beyond the scope of this work, we posit that they can be effectively modelled by adding a spatial dependence to reaction rates or diffusion coefficients. Consequently, we investigate the effect of spatially dependent rates in generic reaction-diffusion processes.

The main motivation comes from the recent study of the critical properties of the diffusive epidemic process (DEP) in the presence of spatial quenched disorder [58]. Critical properties are governed by an exotic fixed point called infinite-disorder fixed point [65] and importantly is surrounded by a Griffiths phase [66]. Griffiths phases originate from the presence of rare regions [67, 68], namely spatial domains with atypical local parameters that can remain locally active for a long time even when the bulk system is inactive. As a consequence, dynamical observables exhibit anomalous scaling and singular behavior over an extended parameter range, rather than only at criticality [66, 68]. In a biological context, and particularly here for cell polarity, time plays an important role shaping the life cycle of the cell [13]. Hence, given the close similarities between the DEP and the NDP model used to model cell polarity [57, 59], a natural question is whether quenched disorder also strongly affects the dynamical properties of the NDP class of models.

To address this question, we start with the simplest analytically tractable case, namely the RR model, where the total number of membrane-bound particles *N*_*m*_ is fixed. We focus on the effect of increasing the contrast between the unbinding rates of the two regions. An analogous analysis can be performed for spatial variations of the recruitment rate. The chosen observable of interest is the mean time during which the majority of membrane-bound particles occupies a given region.

More precisely, we define region *i* as active when the number of particles *n*_*i*_ first exceed a first threshold *n*_*i*_(*t*_1_) ≥ *N*_*th*,1_ up to a second threshold *n*_*i*_(*t*_2_) *< N*_*th*,2_. We then define the active time *T* as *T* = *t*_2_ − *t*_1_. In the RR model with *k* = 2, this definition is equivalent to a mean first-passage problem for the variable *n*_1_ (or equivalently for *n*_2_). We consider trajectories starting at *x* = *N*_*th*,1_, with a reflecting boundary at *b* = *N*_*m*_ and an absorbing boundary at *a* = *N*_*th*,2_. For a birth-and-death process, the mean first-passage time *T* is given by [61]

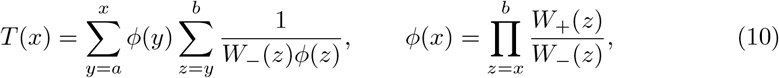

where *W*_+_ and *W*_−_ are defined in Eq. (7). Unless stated otherwise, the mean first-passage times reported for the RR model are obtained by numerically evaluating Eq. (10).

In the following, we restrict ourselves to the case where only the unbinding rates *τ*_*i*_ are heterogeneous and then fix *λ*_*i*_ = *λ, k*_*on,i*_ = *k*_*on*_ ∀*i*, which allows for simpler analytical expressions while preserving the qualitative behavior. The complementary case of heterogeneous recruitment rates leads to analogous results. In Appendix S1 Mean active time in the RR model, we show that in the strongly asymmetric limit *τ*_2_*/τ*_1_ ≫ 1, the mean active time of region 1 scales as

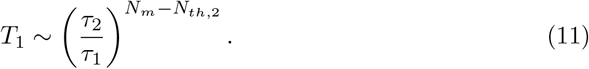

Thus, the mean active time follows a power law in the ratio of unbinding rates, with an exponent proportional to the number of membrane-bound particles *N*_*m*_.

Conversely, in the same limit *τ*_2_*/τ*_1_ ≫ 1, the mean active time of region 2 asymptotically decays as

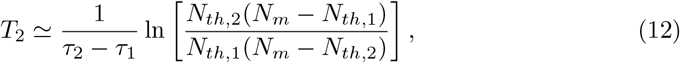

which is independent of *N*_*m*_ if *N*_*th*,1_ and *N*_*th*,2_ are fraction of *N*_*m*_. As a result, a large difference between unbinding rates can lead to a dramatic separation of active times even when *N*_*m*_ is small.

So far, the analysis has assumed a fixed number of membrane-bound particles. However, as discussed in the context of the RCR model, the steady-state deterministic number of cytosolic particles is bounded by *n*_*c*_ *<* min(*τ*_*i*_*/λ*_*i*_). To leading order, this yields

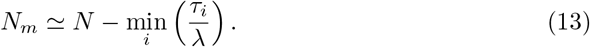

In the asymmetric case of interest, this gives *N*_*m*_ ≃ *N* − *τ*_1_*/λ*, implying that decreasing *τ*_1_ not only favors region 1 dynamically, but also increases the total number of particles bound to the membrane. Quenched disorder therefore has a double effect: it stabilizes one region and simultaneously enhances membrane occupancy.

To compare the RR model with the RCR and NDP models and to define the activation and inactivation thresholds we use the approximation *N*_*m*_ = *N* − min(*τ*_*i*_*/λ*). We set *N*_*th*,1_ = 0.9*N*_*m*_ and *N*_*th*,2_ = 0.1*N*_*m*_. The resulting mean active times are shown in Fig. 2. In all models, the mean active time of the favored region (*τ*_1_ *< τ*_2_) increases rapidly as *τ*_1_ decreases. A difference of only 10% between the unbinding rates already leads to more than five decades of separation between *T*_1_ and *T*_2_. In the RR model with fixed *N*_*m*_, the predicted power-law behavior is clearly visible only at sufficiently small *τ*_1_ (see Appendix S1 Mean active time in the RR model). The difference between fixed and variable *N*_*m*_ highlights the additional contribution of disorder through the increase of membrane-bound particles.

**Fig 2.**
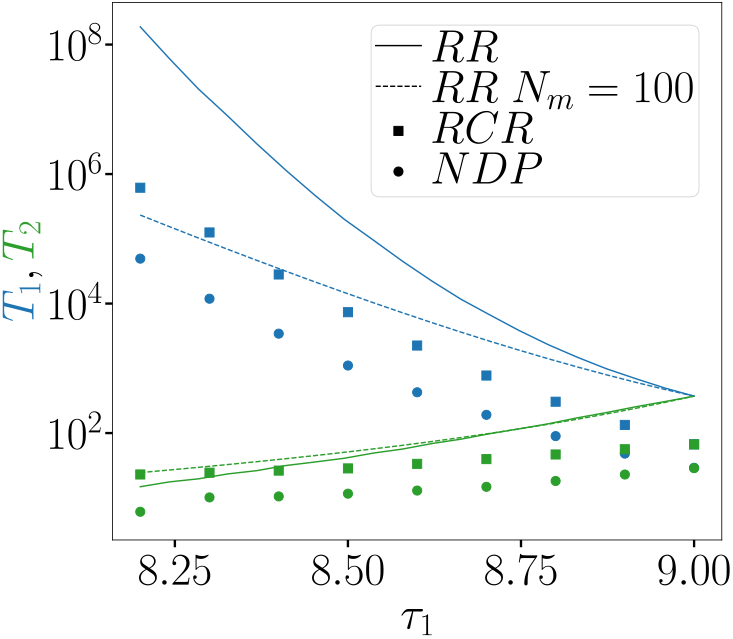
Mean active times of region 1 (*T*_1_, blue) and region 2 (*T*_2_, green) as a function of unbinding rate in region 1, *τ*_1_, with unbinding rate in region 2, *τ*_2_ = 9 and total particle number *N* = 1000. A small asymmetry between *τ*_1_ and *τ*_2_ strongly biases polarization toward the favored region, leading to a large separation between *T*_1_ and *T*_2_. Stochastic simulation results are shown for the NDP (circles) and RCR (squares) models, with analytical estimation for the RR (plain line) model. For the RR model, the dotted line corresponds to a fixed membrane population *N*_*m*_ = 100, whereas the solid line uses *N*_*m*_ = *N* − *τ*_1_*/λ* to account for the increase of the number of membrane-bound particle as *τ*_1_ decreases.

A systematic reduction of *T*_1_ is observed when going from the RR to the RCR and NDP models. The discrepancy between RR and RCR originates from fluctuations in *N*_*m*_, which can destabilize active regions when the membrane population becomes temporarily small. The remaining difference between RCR and NDP reflects finite-size and diffusion effects, controlled by the system size *L* or equivalently the membrane diffusion coefficient *D*. Increasing *L* or decreasing *D* leads to convergence of the NDP results toward those of the RCR model.

We now fix all rates and vary only the total number of particles *N*, or equivalently the expected membrane population *N*_*m*_. For *τ*_1_ *< τ*_2_, the exponential scaling of *T*_1_ with *N*_*m*_ predicted by Eq. (11) remains valid when *N*_*m*_ ≫ 1 as detailed in Appendix S1 Mean active time in the RR model. To highlight the role of disorder, we compare the results with the homogeneous case given by *τ*_1_ = *τ*_2_ and *k*_*on*,1_ = *k*_*on*,2_. In Appendix S1 Mean active time in the RR model, we show that in this case the mean active time scale asymptotically as

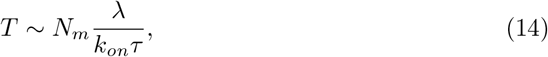

provided that the thresholds *N*_*th*,1_ and *N*_*th*,2_ are fraction of *N*_*m*_. Hence, the mean active time grows linearly with *N*_*m*_ in the homogeneous case, in sharp contrast with the exponential growth observed when *τ*_1_ ≠ *τ*_2_. Fixing *N*_*th*,1_ = 0.9*N*_*m*_ and *N*_*th*,2_ = 0.1*N*_*m*_, Fig. 3 shows the dependence of *T*_1_ and *T*_2_ on *N*_*m*_ for both cases and for the three models. The mean active times *T*_1,2_ in the NDP and RCR show a similar grow, linear and exponential, as for the RR model with *N*_*m*_ for, resp. *τ*_1_ = *τ*_2_ and *τ*_1_ *< τ*_2_.

**Fig 3.**
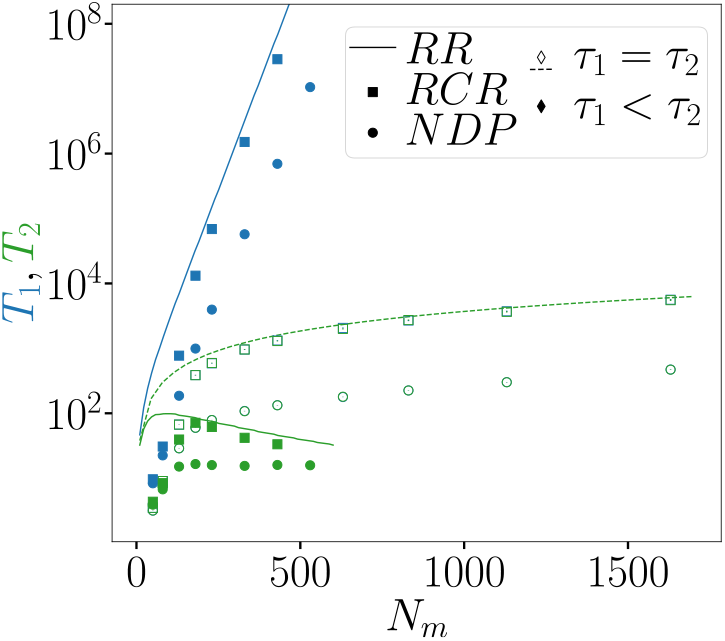
Mean active times for the polarization of region 1 (*T*_1_, blue) and region 2 (*T*_2_, green) as a function of the estimated number of membrane-bound particles, *N*_*m*_ = *N* − *τ*_1_*/λ*. Data are shown for the homogeneous case (*τ*_1_ = *τ*_2_ = 9; open symbols and dotted lines, where *T*_1_ = *T*_2_) and in the disordered regime (*τ*_1_ = 8.7, *τ*_2_ = 9; filled symbols and solid lines). Spatial heterogeneity significantly alters the scaling with *N*_*m*_. Symbols represent the NDP (circles) and RCR (squares), while lines denote the RR model. For the latter, the large-*N*_*m*_ asymptotic behavior of *T*_1_ is in agreement with Eq. (11) for the homogeneous case and Eq. (14) for the disordered case.

Strikingly, quenched disorder has a strong impact on measurable dynamical observables such as active times. Even weak disorder produces dramatic effects when the number of particles is large, and similarly for a stronger disorder with a smaller number of particles. From a biological perspective, this suggests that locally tuning reaction rates in systems with positive feedback provides an efficient mechanism to generate robust and localized signaling at low cost. Switching back the rates to the bulk value will either make the polarized region to diffuse in the whole system as no region is no longer favored or remove the polarization region because of the change of the polarization threshold. Such effects have been observed experimentally in mating fission yeast cells [64].

Although we have focused on two regions of equal size, the conclusions remain valid for unequal regions, which mainly affect the effective binding rates *k*_*on,i*_. Membrane diffusion, however, imposes a minimal region size set by the competition between recruitment and escape times, and always weakens localization. These results naturally generalize to systems with multiple regions and heterogeneous parameters.

### Emergence of oscillations

In this section, we investigate the emergence of oscillations induced by quenched spatial disorder. Specifically, we consider a system composed of four spatial regions, arranged as two pairs of identical regions, as schematized in Fig. 4(a). Such a geometry can represent a rod-shaped cell, where the physical and biochemical properties at the cell tips differ from those along the long axis. This setup is motivated by the oscillatory behaviors observed in many rod-shaped cells, including fission yeast [11, 50, 51] and *Escherichia coli* [69–71].

**Fig 4.**
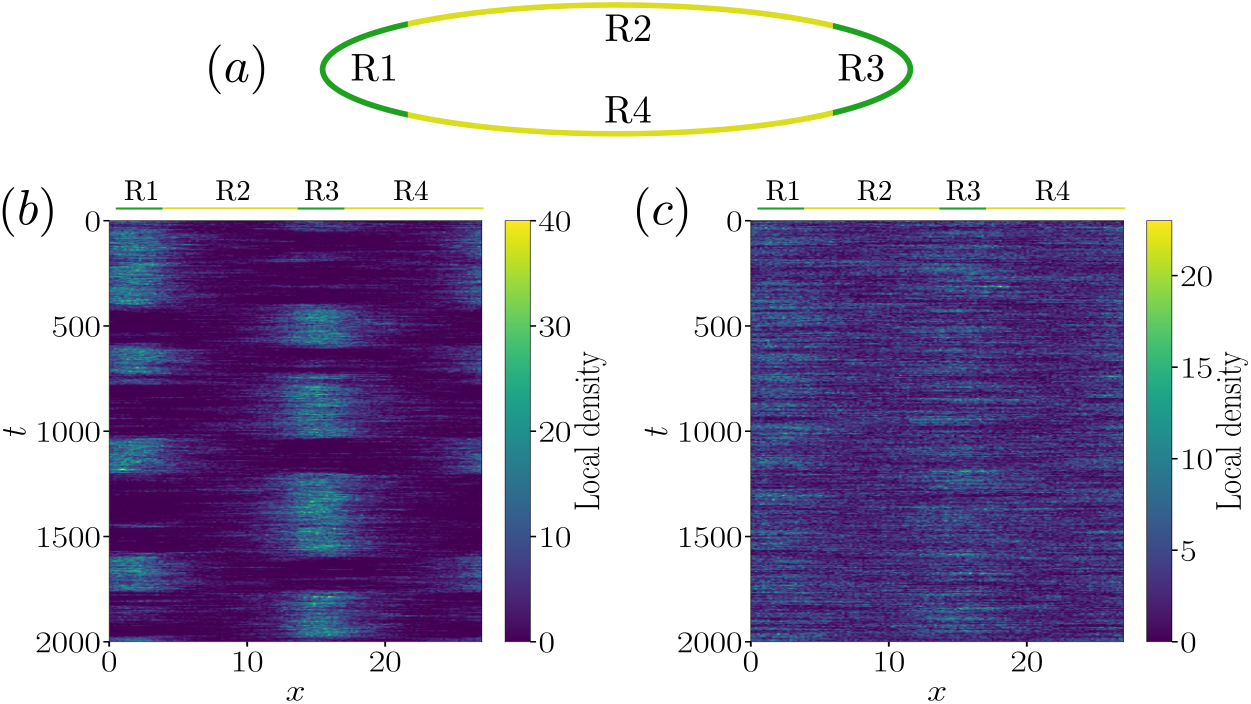
(a) Quenched-disorder setup mimicking an elongated cell. By symmetry, regions R1 and R3 share identical parameters, as do regions R2 and R4. (b) Kymograph of the density of membrane-bound particles in the NDP model using the geometry shown in panel (a). Oscillations between the two tips (R1 and R3) emerge only in the presence of both positive feedback and quenched disorder, implemented through spatially dependent unbinding rates *τ*_1_ = *τ*_3_ = 8.5 in R1 and R3 and *τ*_2_ = *τ*_4_ = 9 in R2 and R4. Due to membrane diffusion (*D* = 1.2), particles are also present in regions R2 and R4. (c) Same configuration but with larger membrane diffusion (*D* = 5), which suppresses the oscillatory behavior.

From a theoretical standpoint, sustained temporal oscillations can arise through several mechanisms, such as negative feedback with at least three components [66, 72], combinations of positive and negative feedback [73, 74], explicit time delays [75], or sufficiently strong nonlinearities [76]. In spatially extended systems, additional mechanisms become available. Notable examples include Turing instabilities [14], traveling waves [77], activator–inhibitor dynamics [78, 79], biased diffusion favoring aggregation [53], and delayed feedback in space [50]. Here, we show that quenched disorder alone, in a mass-conserved stochastic system with positive feedback, is sufficient to generate stochastic switching.

We focus on the same class of models introduced in Section Models. As for spontaneous polarization, the oscillatory behavior in fission yeast is primarily driven by the GTPase Cdc42 [11, 50]. Most theoretical descriptions of Cdc42 oscillations reduce the system to two compartments corresponding to the cell tips, coupled by diffusion [43–46, 48]. This simplification is motivated by experimental observations showing that high Cdc42 concentration is predominantly localized at the poles [50].

Here, instead, we postulate that intrinsic spatial heterogeneities exist prior to any reaction dynamics. In particular, we assume that the cell tips (regions R1 and R3) are intrinsically favored relative to the lateral regions (R2 and R4). This assumption is motivated by two main considerations. First, microtubules along the long axis deliver proteins such as Tea1, Tea4, and Pom1, which promote Cdc42 localization at the poles by excluding Cdc42 GAPs and recruiting Cdc42 GEFs [80–83]. Rather than modeling these molecular details explicitly, we incorporate their net effect as a locally reduced unbinding rate *τ* at the poles. An equivalent phenomenology would be obtained by locally increasing the recruitment rate *λ*. More generally, other kinds of spatial cues could be similarly taken into account. Second, recent measurements indicate that the diffusion of some cytosolic proteins is slower near the poles than at the cell center [84]. More generally, both numerical and analytical studies have emphasized the influence of geometry on polarization [27, 28, 85] and pattern formation [86]. In the DEP framework, heterogeneities in diffusion rates affect critical properties in a manner analogous to disorder in *τ* or *λ* [58]. For clarity, we therefore focus on disorder in the unbinding rate, noting that similar conclusions hold for disorder in recruitment or diffusion rates.

As shown in the previous section, even a weak local advantage—such as a reduced unbinding rate or enhanced recruitment—can produce large differences in dynamical observables. In the rod-like geometry considered here, favoring the poles implies that membrane dynamics along the long axis (regions R2 and R4) becomes negligible. This provides an alternative justification for models that focus exclusively on tip dynamics [43–46, 48]: not because reaction rules differ elsewhere, but because the relative difference of rates implies a very unlikely and short polarization away from the poles. This observation also justifies the use of the simplified models introduced previously, RCR and RR, with two identical active regions corresponding to R1 and R3, where only the spontaneous binding rate needs to be rescaled according to region size, *k*_*on,i*_ = *k*_*on*_*L*_*i*_*/L*.

Within this interpretation, the sum of the mean active times of regions R1 and R3, as given by Eq. (14), naturally defines the oscillation period between the two poles. Importantly, these oscillations are not periodic in the deterministic sense, but corresponds to a stochastic switching.

To demonstrate the emergence of oscillations in the NDP model, we choose *L*_1_ = *L*_3_ = 3.5 *µ*m and *L*_2_ = *L*_4_ = 10 *µ*m, consistent with the geometry of *Schizosaccharomyces pombe* [54, 87]. *All other parameters are identical to those of the previous section except for the diffusion D* along the membrane whose effect is studied in the following. Fig. 4(b) shows a representative kymograph obtained for *N* = 1400, *D* = 1.2, and *τ*_1_ = *τ*_3_ = 8.5, corresponding to a weak disorder favoring the poles. High densities of membrane-bound particles are localized predominantly in regions R1 and R3, with some spreading near their boundaries due to diffusion. Notably, simultaneous strong activation of both poles is never observed, resulting in an irregular oscillatory pattern.

Nevertheless, diffusion along the membrane in the NDP model introduces a new mechanism absent from the RCR and RR models: particles can migrate directly from one pole to the other while undergoing recruitment and unbinding. In Fig. 4(c), the kymograph obtained with *D* = 5 no longer have oscillations, instead the system is homogeneous albeit with still a larger density at the poles. This effect can be incorporated phenomenologically in the RR model by adding a direct exchange flux between the two regions. The transition rates in Eq. (7) then become

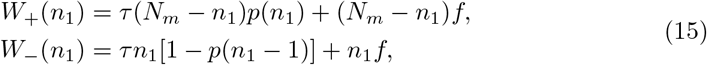

where *f >* 0 quantifies the strength of direct exchange and we have considered identical unbinding rate *τ* in both regions as the dynamic between R1 and R3 is studied.

To characterize the resulting dynamics, we define *P*_1_ as the probability that a single pole is strongly activated corresponding to *n*_1_ *< N*_*m*_*/*4 or *n*_1_ *>* 3*N*_*m*_*/*4. The complementary probability 1 − *P*_1_ is interpreted as both poles being simultaneously active. Using Eq. (9), *P*_1_ is readily computed. Its dependence on *f* at fixed *N*_*m*_ and on *N*_*m*_ at fixed *f* is shown in Fig. 5(a). For small *f* or with few particles *N*_*m*_, *P*_1_ is close to 1, corresponding to the exclusive activation of a single pole. Increasing *f* or *N*_*m*_ induces a smooth crossover toward a regime dominated by direct exchange, where both poles are active. An estimate of the crossover point *P*_1_ = 0.5 can be obtained by noting that a flat stationary distribution implies *W*_+_(0) = *W*_−_(1), yielding 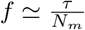. This estimate agrees well with the numerical results shown in Fig. 5.

**Fig 5.**
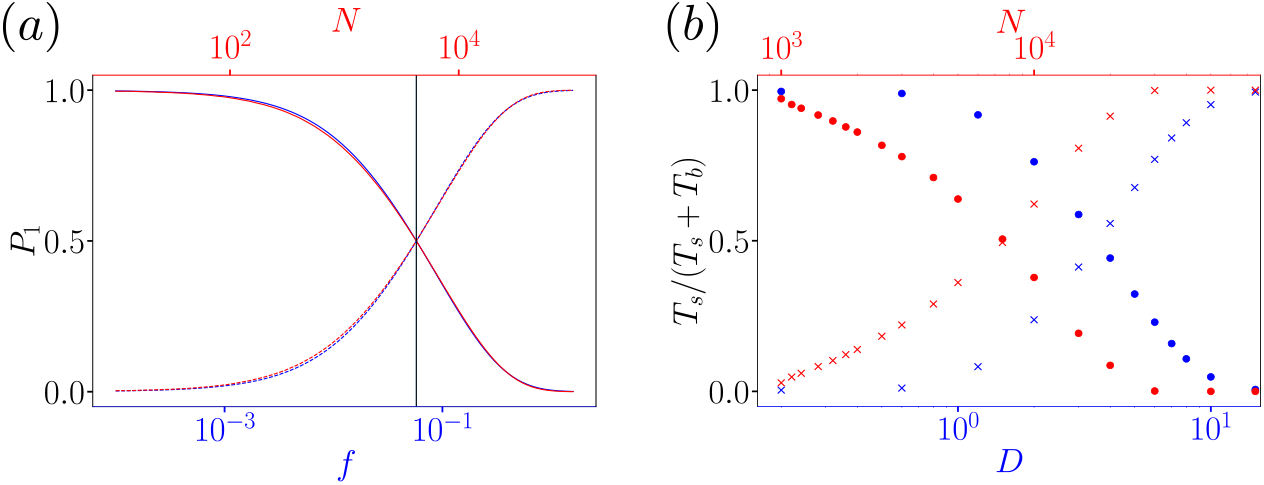
(a) The RR model with an additional exchange rate *f*. Probability *P*_1_ of observing a single activated pole (solid lines) as a function of the exchange rate *f* at fixed membrane population *N*_*m*_ = 150 (blue) and as a function of *N*_*m*_ at fixed *f* = 0.002 (red). Dotted lines show the complementary probability 1 − *P*_1_. The vertical black line indicates the estimate position *f* = *τ/N*_*m*_ of the transition. (b) The NDP model with two favored regions (*τ*_1_ = *τ*_3_ = 8.5). Fraction of activity spent in the single-pole state, *T*_*s*_*/*(*T*_*s*_ + *T*_*b*_) (circles), and in the two-pole state, *T*_*b*_*/*(*T*_*s*_ + *T*_*b*_) (crosses), where *T*_*b*_ denotes the time during which both poles are active. The transition between these regimes is controlled by the direct particle flux exchanged between the two favored poles, which promotes global homogenization when the two-pole state dominates. Blue symbols show the dependence on the diffusion coefficient *D* at fixed *N* = 1400, while red symbols show the dependence on the total particle number *N* at fixed *D* = 1.2.

To identify the same transition in the NDP model, new observables are required due to the finite occupancy of regions different from the poles, i.e., R2 and R4. We therefore define the pole activity based on the relative imbalance between the two tips. A single pole is set active when it contains at least nine times more particles than the opposite pole, and is set inactive when the populations at the two poles are equal. Both poles are set active when |*n*_1_ − *n*_3_ |*/* max(*n*_1_, *n*_3_) *<* 0.2, and set inactive when this ratio exceeds 0.6. Although these thresholds are arbitrary, they capture qualitative trends. We denote by *T*_*s*_ the cumulative time during which only one pole is active, and by *T*_*b*_ the time during which both poles are active.

Fig. 5(b) shows the ratio *T*_*s*_*/*(*T*_*s*_ + *T*_*b*_) as a function of the total particle number *N* at fixed diffusion, and as a function of the diffusion coefficient *D* at fixed *N*. The results closely mirror those of the RR model with exchange: a smooth crossover from a monopolar regime to a bipolar regime is observed as *N* or *D* increases.

In the NDP model, however, simultaneous activation of both poles necessarily coincides with near-homogenization of the entire membrane, as seen in Fig. 4(c) at large *D*. Consequently, two active poles cannot coexist while the rest of the membrane remains inactive. This homogenization is enabled solely by membrane diffusion. In its absence, i.e., *D* = 0, the density at each pole would grow without bound with *N*, and the mean active time would increase linearly with the total particle number as expected from Eq.14.

### Coexistence

The potential localised infinite density at the membrane exposes a fundamental breakdown in the approximation of a well-mixed cytosol inherent to the NDP model. When membrane-bound particle densities are large, reaction timescales can become faster than the time required for cytosolic diffusion to homogenize local concentrations, thereby invalidating the assumption of spatial uniformity. To investigate this regime, we adopt the DEP description, which explicitly resolves cytosolic transport by introducing a cytosolic diffusion coefficient *D*_*c*_ ≫ *D*_*m*_. In this discrete framework, we use the subscript *m* for the membrane transport parameter *D*_*m*_ to distinguish it from *D*_*c*_. Note that unlike the continuum coefficient *D* (in units of µm^2^ min^−1^), the discrete parameter *D*_*m*_ represents a hopping rate with units of frequency (min^−1^). Technical details regarding the spatial discretization are provided in Appendix S3 Numerical simulations.

We adopt the same definitions for the cumulative duration of single-pole activity, *T*_*s*_, and simultaneous two-pole activity, *T*_*b*_. In Fig. 6(a), we plot the fraction of time spent in the monopolar state, defined as the ratio *T*_*s*_*/*(*T*_*s*_ + *T*_*b*_), as a function of the total number of particles *N*. Similar to the NDP model, the DEP exhibits a transition from a monopolar to a bipolar regime. However, two distinct features emerge in the presence of explicit cytoplasmic diffusion. First, the transition is significantly sharper in the DEP model: it occurs over a narrow range of particle numbers (*N* ≈ 1000–2000), whereas in the NDP framework, the crossover extends over more than an order of magnitude in *N* (Fig. 5(b)). Second, both the threshold and the width of the transition depend strongly on the cytosolic diffusion coefficient *D*_*c*_. Notably, as *D*_*c*_ increases, the DEP results (red circle) progressively converge toward the NDP limit (green circle).

**Fig 6.**
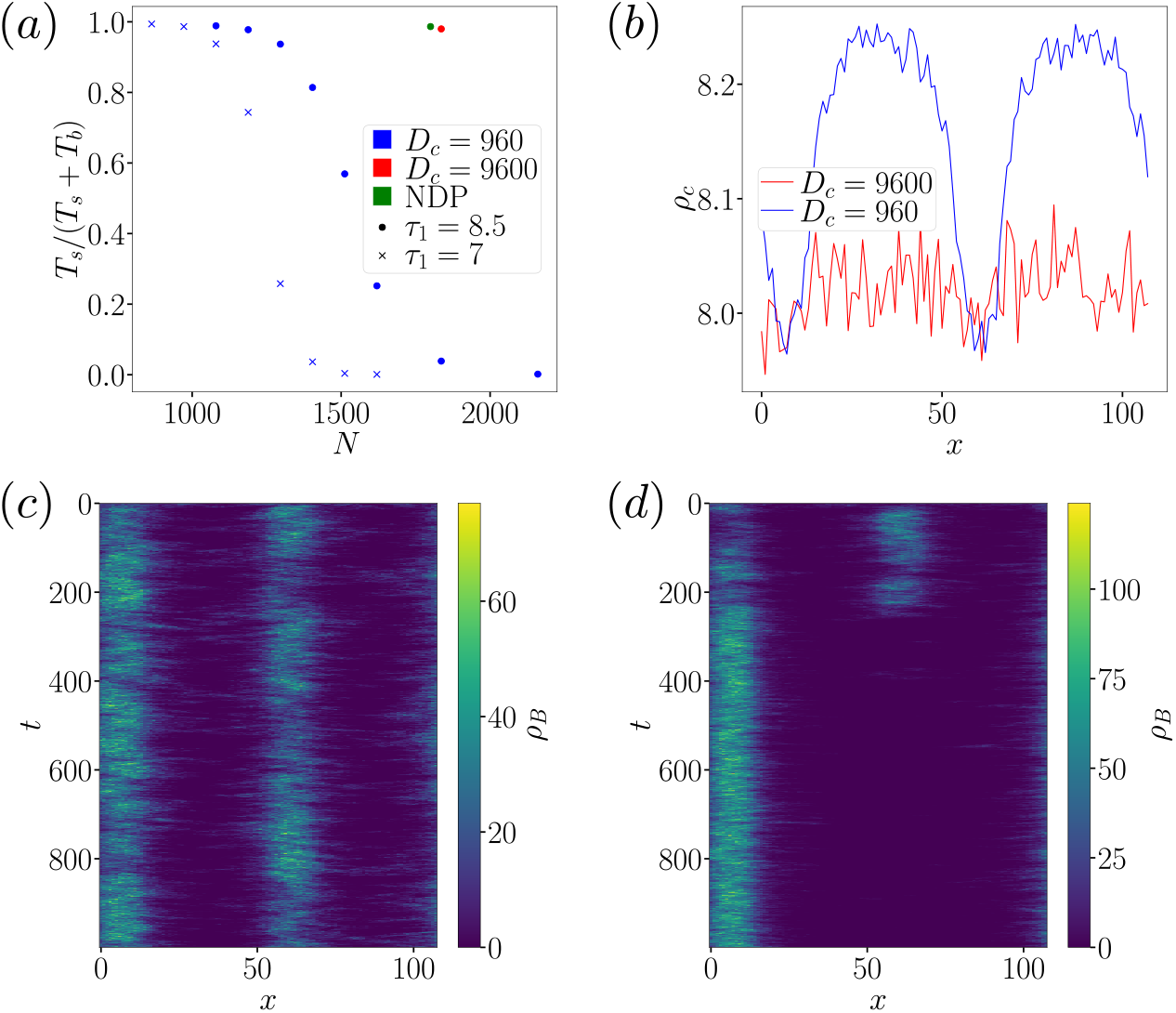
The DEP displays the transition between a monopolar and a bipolar state. (a) Fraction of activity in the single-pole state, *T*_*s*_*/*(*T*_*s*_ + *T*_*b*_), as a function of the total particle number *N* = *ρL* for *D*_*c*_ = 960 (blue), *D*_*c*_ = 9600 (red), and in the NDP limit (green). Symbols indicate *τ*_1_ = *τ*_3_ = 8.5 (circles) and *τ*_1_ = *τ*_3_ = 7 (crosses). (b) Spatial profiles of the cytosolic particle density *ρ*_*c*_(*x*) for *D*_*c*_ = 960 and *D*_*c*_ = 9600 at *ρ* = 17 and *τ*_1_ = *τ*_3_ = 8.5. (c,d) Representative kymographs associated for *D*_*c*_ = 960 (c) and *D*_*c*_ = 9600 (d). Two poles can remain active when the cytosolic density becomes sufficiently inhomogeneous, effectively limiting the occupation of a single pole. Increasing *D*_*c*_ restores a winner-takes-all mechanism.

To elucidate the influence of cytosolic diffusion, Fig. 6(c) displays kymographs for *D*_*c*_ = 960 and *N* = 1800. In this regime, stable bipolarity is observed even while the particle density remains low in the inter-polar region. This behavior stands in sharp contrast to the NDP model, where a single pole persists as long as the background density is low, and simultaneous activation of both poles emerges only once the system becomes nearly homogeneous (see Fig. 4(c)). As *D*_*c*_ increases, the DEP dynamics converge toward this NDP phenomenology. For instance, at *D*_*c*_ = 9600 (Fig. 6(d)), while two poles can transiently coexist, this state is significantly less stable, exhibiting frequent reversions to a monopolar configuration.

The discrepancy between the DEP regimes (moderate vs. large *D*_*c*_) and the NDP model stems from the spatial heterogeneity of the cytosolic density *ρ*_*c*_(*x*). Fig. 6(b) displays the stationary cytosolic profiles. At *D*_*c*_ = 960, the distribution is markedly inhomogeneous, exhibiting pronounced depletion within the favored regions R1 and R3. Conversely, at *D*_*c*_ = 9600, the density approaches homogeneity, although residual depressions in R1 and R3 remain discernible.

This behavior can be rationalized via a steady-state flux balance argument. Consider the interface between a domain with high membrane density 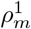 and a domain with negligible membrane density. Requiring the total diffusive flux (cytosolic plus membrane-bound) to be uniform yields the condition:

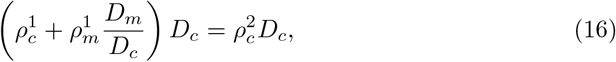

where 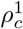 and 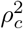 denote the cytosolic densities in the respective regions. From Eq. (16), it follows that a nearly homogeneous cytosol 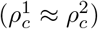 requires the transport to be dominated by cytosolic diffusion, specifically 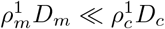. This criterion explains the progressive homogenization of *ρ*_*c*_(*x*) observed in Fig. 6(b) as *D*_*c*_ is increased.

This flux-balance mechanism elucidates the conditions required for stable bipolarity. Consider a system with a single favored region (R1). If this region is favored, the local density of membrane-bound particles, 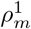, is large, creating a significant sink that depletes the local cytosolic density, 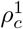. However, due to the finite diffusivity *D*_*c*_, this depletion zone remains spatially localized. Consequently, provided the cytosolic density in a distant region (e.g.,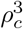) remains above the critical activation threshold, a second pole can emerge and persist. The stability of this coexistence depends on the density difference 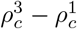 before the emergence of the second polarized pole.

This reasoning also rationalizes the counter-intuitive observation that increasing the affinity of the polar regions (R1 and R3) relative to the lateral regions (R2 and R4) actually promotes simultaneous activation. This effect is evident in Fig. 6(a): the transition from a monopolar to a bipolar state shifts to lower total particle numbers *N* when the spontaneous unbinding rates in the favored regions (*τ*_1_ = *τ*_3_) are reduced. It arises for two reasons. First, reducing these rates lowers the critical particle number required to sustain polarization. Second, because *D*_*c*_ is finite, the local increase of membrane-bound particles in one region does not necessary result in a commensurate global reduction of the cytosolic density. Thus, increasing the affinity lowers the activation threshold without proportionally increasing the global competition, thereby facilitating the onset of bipolarity at lower *N*. This behavior stands in marked contrast to the NDP, RCR, and RR models, which are characterized by a globally well-mixed cytosol. In those models, increasing the affinity of favored regions enhances the direct, instantaneous competition for the shared cytosolic pool by increasing the mean active time of a single polarized region, see Eq.14.

### Cell growth

A change in the number of polarized regions during the cell cycle is well documented in living systems. In fission yeast, cells initially grow in a monopolar manner, with the polarity regulator Cdc42 concentrated at a single tip. Above a characteristic cell length, growth switches to a bipolar mode in which both ends become active [50, 54]. This transition, known as New End Take-Off (NETO), has been attributed to the increase in available polarity regulators and to saturation mechanisms that stabilize bipolar configurations [44, 50]. These observations suggest that geometric growth and resource availability can qualitatively reshape polarization dynamics.

A qualitatively similar scenario arises in the DEP model with quenched disorder. Starting from a single active pole, as the system elongates along its long axis (by adding new sites in regions R2 and R4), the spatial distribution of cytoplasmic particles becomes progressively inhomogeneous. In particular, the background density *ρ*_*c*_(*x*) increases away from the initially active pole. When this density exceeds a threshold, the system can sustain simultaneous activation of both tips, leading to coexistence of polarized regions.

To model cell growth, we increase the system size incrementally by inserting lattice sites at the centers of regions R2 and R4. Every time interval Δ*t* = 20, one site occupied by ⌊*ρ* ⌋ (or ⌊*ρ* ⌋+ 1 to maintain the prescribed mean density *ρ*) cytoplasmic particles is added symmetrically, up to a final time *t* = 1080, corresponding to a doubling of the cell length. Compared to previous sections, we use a larger spontaneous attachment rate *k*_on_ = 0.02 to accelerate the competition, as growth speed introduces an additional timescale; keeping the smaller *k*_on_ used previously would have not make a clear change during the growth.

We first consider equally favored tips, *τ*_1_ = *τ*_3_ = 8.2. A representative kymograph at *ρ* = 10 is shown in Fig. 7(a). At early times, when the system is short, a single pole dominates and stochastic switching between the two tips is observed. As the cell elongates, configurations in which both poles are simultaneously active become increasingly frequent, while single-pole states are progressively suppressed. As done in the previous section, a single active pole can be detected by the ratio

**Fig 7.**
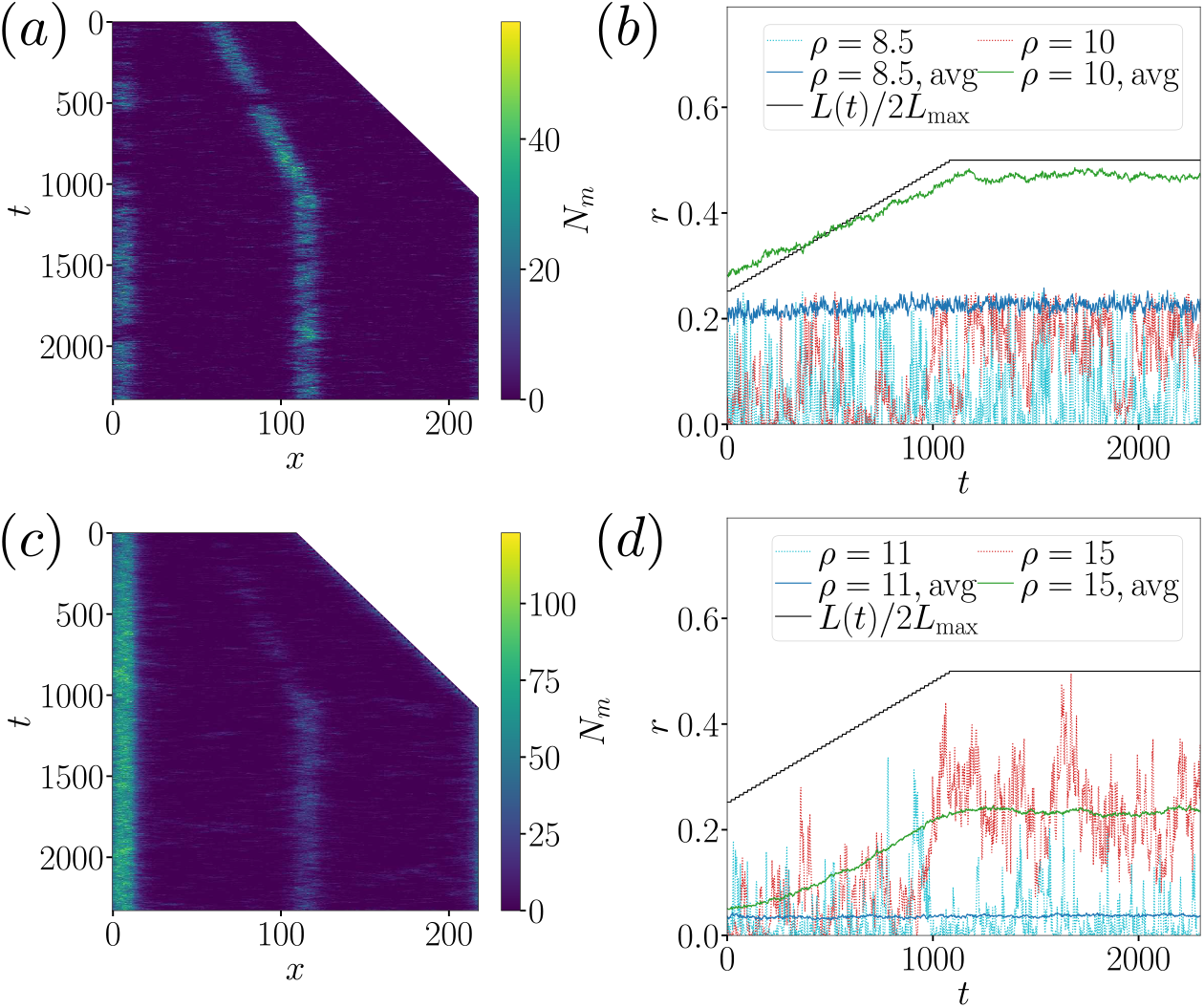
Effect of spatial heterogeneity on polarization dynamics during cell growth in the DEP. (a,b) Symmetric case: regions R1 and R3 are identical (*τ*_1_ = *τ*_3_ = 8.2). (a) Kymograph of the membrane-bound particle density *N*_*m*_ in a cell growing at constant density *ρ* = 10 by insertion of new sites at mid-cell (regions R2 and R4). (b) Time evolution of the ratio *r* = min(*n*_1_, *n*_3_)*/* max(*n*_1_, *n*_3_), where *n*_1_ and *n*_3_ denote the total number of membrane-bound particles in R1 and R3. Results are shown for *ρ* = 10 (green) and *ρ* = 8.5 (blue), averaged over at least 700 realizations; dotted lines show representative trajectories (rescaled by a factor 1*/*4 for clarity). Small values of *r* correspond to single-pole dominance, whereas larger values indicate coexistence of two poles. Cell growth promotes stable bipolar activation for *ρ* = 10, while stochastic switching persists for *ρ* = 8.5, which therefore serves as a reference case without coexistence. During growth the cell length increases from *L*_0_ = 108 to *L*_max_ = 216 (black line). (c,d) Asymmetric case: region R1 is more favorable than R3 (*τ*_1_ = 8, *τ*_3_ = 8.5). (c) Representative kymograph at *ρ* = 15. (d) Time evolution of the same ratio *r*; dotted lines show representative trajectories. In this regime, cell growth promotes activation of the less favored pole, reminiscent of the NETO transition. Other parameters: *D*_*c*_ = 960, *D*_*m*_ = 9.6, *τ*_2_ = *τ*_4_ = 9, *λ* = 1.08, and *k*_on_ = 0.02.

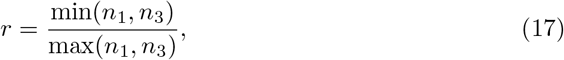

where *n*_1_ and *n*_3_ denote the total number of membrane-bound particles in R1 and R3. Small values of *r* correspond to single-pole dominance, whereas *r* ≈ 1 indicates perfect equalization, which, in practice can not be reached due to stochastic fluctuations. Similarly, because stochastic switching prevents *r* from vanishing identically even in a monopolar regime, we include *ρ* = 8.5 as a reference case where coexistence does not occur. In this regime, typical trajectories repeatedly return to *r* ≈ 0, and the ensemble-averaged (over at least 700 independent samples) *r* remains constant throughout growth. By contrast, for *ρ* = 10, the averaged ratio increases with *L*(*t*), indicating that an increasing fraction of time is spent in configurations where both poles are comparably populated, as shown in Fig. 7(b). Hence, that cell growth promotes coexistence.

However, in fission yeast, NETO occurs without stochastic switching prior to bipolar growth. To reproduce a closer phenomenology within our model, we introduce an intrinsic asymmetry by setting *τ*_1_ = 8 and *τ*_3_ = 8.5, making R1 more favorable. A representative kymograph at *ρ* = 15 is shown in Fig. 7(c). Initially, only the favored pole (R1) is active. As the cell elongates and cytoplasmic density being increasingly inhomogeneous, a second polarized region emerges at R3 and eventually coexists with the dominant pole, although with a smaller particle number. The corresponding evolution of *r* in Fig. 7(d) confirms this transition: for *ρ* = 15, *r* increases with *L*(*t*), whereas for the reference density *ρ* = 11 it remains small and constant, indicating that the second pole is unstable. Biologically, a stronger initial favorability of the old pole may reflect structural asymmetries, such as pre-existing spatial cue or microtubule organization that facilitate Cdc42 recruitment at the inherited tip [88].

Despite the qualitative parallels with NETO, several limitations should be emphasized. The introduction of quenched disorder — i.e., assuming that some regions are intrinsically more favored — over-simplifies the underlying dynamics. It effectively amounts to treating the rest of the cellular processes as time-invariant (frozen) and absorbing their influence into renormalized parameters of the model. This approximation is clearly incomplete. For instance, microtubules are not stable structures independent of Cdc42 activity, but are dynamically coupled to polarity establishment [88, 89]. Allowing parameters such as the unbinding time *τ* or recruitment rate *λ* to depend on the local Cdc42 density, or explicitly including additional regulatory species, would provide a more realistic description of these feedbacks. Finally, experiments show that Cdc42 activity at the two tips can converge toward comparable intensities as the cell grows [50]. Such convergence would not be possible in the present framework if the two tips remain permanently unequally favored, further highlighting the limitations of modeling pole asymmetry as quenched disorder.

## Discussion

Quenched disorder has been extensively studied in statistical physics [33, 34, 68, 90–92], particularly for its impact on critical phenomena. By contrast, its role appears comparatively underexplored in other disciplines, including biophysics. One possible reason is that many living systems are not finely tuned to a critical point: for conventional disorder with finite mean and variance, and in the absence of frustration, quenched heterogeneities are generally not expected to qualitatively affect the dynamics far from criticality [93]. However, disorder can also induce much more radical behavior. Highly unconventional fixed points, known as infinite-disorder fixed points (IDFPs), were first identified in disordered quantum systems [65] and later in non-equilibrium classical models [94, 95]. These fixed points are accompanied by extended Griffiths phases [39], in which rare but long-lived regions dominate the dynamics and give rise to anomalously slow, power-law relaxation over broad parameter ranges away from criticality. Recently, IDFPs have been identified in the diffusive epidemic process (DEP) [58], a minimal model with positive feedback that shares key ingredients with theoretical descriptions of spontaneous cell polarity [57, 59]. In the context of polarity, time plays a central role: processes such as budding require that a polarized state persist for a sufficiently long duration before downstream events are triggered [13]. Quenched disorder may therefore strongly influence biologically relevant observables, even when the system is not tuned close to a critical point. In this work, we have shown that a minimal mass-conserved stochastic system with positive feedback is highly sensitive to spatial disorder. Regions with only a slight local advantage become exponentially more likely to host polarity than less favored ones. From this perspective, quenched disorder can be viewed as a simple and efficient mechanism for generating robust and spatially localized polarity.

This naturally raises a fundamental question: why and how should quenched disorder arise in a cellular context? At first glance, frozen heterogeneities may seem implausible at the scale of a cell, where many processes occur rapidly compared to the timescales typically associated with quenched disorder in condensed-matter systems. However, the notion of “quenched” disorder must always be understood relative to a relevant timescale. If the biological question concerns, for instance, the initiation of budding, then any spatial heterogeneity that persists over the timescale of bud formation effectively acts as quenched disorder.

Beyond this conceptual point, quenched disorder also provides a useful modeling abstraction. Spatial heterogeneities in cells can originate from a wide range of sources, including geometry [26–29], microtubules [96], actin microfilaments [97], ion gradients [97], morphogenetic proteins [86, 98, 99], cytoplasmic heterogeneities [100–102], or external chemical gradients [25, 64]. Explicitly modeling all these effects is computationally costly; using quenched disorder as an effective description can therefore serve as a powerful first step to explore possible dynamical outcomes before pursuing more detailed and specific models.

Beyond its practical utility, the success of a description based on quenched disorder suggests that interactions other than those generating the heterogeneities may be subdominant for the phenomena of interest. In this sense, simplifying a system through quenched disorder can provide genuine conceptual insight. For example, we have shown that oscillatory behavior can emerge in a two-species, mass-conserved stochastic system with a simple positive feedback loop subject to spatial disorder. This stands in contrast to many existing models of spatial or temporal oscillations, which typically rely on more elaborate ingredients such as delayed feedback [50], combined positive and negative feedback loops [52], activator–inhibitor mechanisms [78, 79], or biased diffusion [53]. That said, the oscillations observed here arise from stochastic switching and are therefore not strictly periodic. This suggests that additional mechanisms—such as explicit delays or negative feedback—are required to reproduce the more regular oscillatory dynamics reported in experiments.

A closely related issue concerns the fate of multiple polarity sites. In some biological systems, competition between polarity zones leads to the survival of a single site [9, 103, 104], whereas in others several sites can coexist [11, 105], with possible transitions between single- and multi-peak states [54, 106, 107]. Winner-takes-all scenarios based on positive feedback and mass conservation have been extensively studied to account for the selection of a single dominant polarity spot [21, 22, 103]. Nevertheless, when the biological timescale of interest is shorter than the competition timescale, multiple, typically unequal, polarity sites may persist for extended periods [103]. Several alternative mechanisms have been proposed to stabilize the coexistence of multiple polarity sites, including negative feedback loops [108, 109], advective transport or convection [110], saturation effects [55, 56], explicit production and degradation terms [108, 111], or activator–substrate depletion mechanisms [24, 112]. Here, within the same minimal mass-conserved framework, we show that coexistence can also arise from a different and simple mechanism: a non-homogeneous spatial distribution of cytoplasmic particles induced by finite diffusion, which is not fast enough to homogenize the system relative to the local dynamics within polarity sites. This behavior emerges without introducing additional species or feedback loops; instead, the effective complexity is encoded in space-dependent rates generated by quenched disorder. While this mechanism remains to be tested experimentally, it captures features observed in living cells, such as the transition from a single active pole to the coexistence of two active poles as the cell grows [50] or the emergence of multiple polarized regions as the total amount of polarity proteins increases [24], with the important caveat that the existence of multiple spatially favored regions cannot be directly assessed in those experiments. The present results therefore suggest another possible interpretation of such observations, rather than a direct explanation, while keeping in mind that a collection of mechanisms (negative feedback, saturation, …), rather than a single one, is likely to be present.

Finally, many homogeneous models have been remarkably successful in explaining experimental observations. However, if these models are, in principle, expected to be sensitive to quenched disorder, their surprising effectiveness itself calls for explanation and may provide valuable insights. Does this success imply that the biological system is effectively homogeneous at the relevant scales? Or do spatial heterogeneities fluctuate rapidly enough to average out over experimental timescales, e.g. correspond to an annealed and not quenched disorder? Alternatively, could it signal the presence of unmodeled processes that actively suppress the effects of quenched disorder?

Although the notion of quenched disorder is not always straightforward to apply to living cells, there are situations in which it can be biologically well motivated. In such cases, this perspective opens the door to applying a broad range of theoretical tools developed over the past decades for the study of disordered systems, potentially yielding new insights into cellular organization and dynamics.

## Acknowledgments

This work was partially supported by grants from the National Science and Technology Council, Taiwan (Grant No. NSTC 111-2112-M-001-027-MY3 and 114-2112-M-001-062) and Academia Sinica Career Development Award (Project No. AS-CDA-114-M02). VA acknowledges support from Academia Sinica Postdoctoral Scholar Program.

## Supporting information

### S1 Mean active time in the RR model

- We derive asymptotic expressions for the mean absorbing times given by Eq. 10 in the limit *τ*_2_ ≫ *τ*_1_.

– **In Region 1**. We first consider the ratio appearing in Eq. 10:

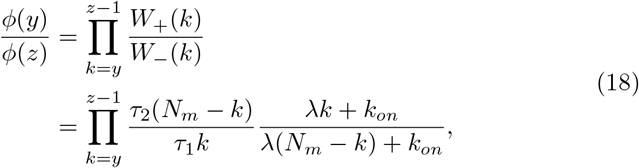

for 0 *< y* ≤ *z* ≤ *N*_*m*_.

We neglect the corrections *k*_*on*_*/*(*λk*) and *k*_*on*_*/*(*λ*(*N*_*m*_ − *k*)) to obtain

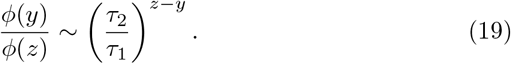

Substituting Eq. 19 into Eq. 10 gives

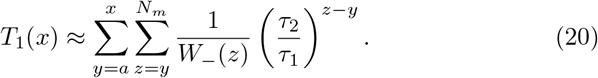

In the limit *τ*_2_ ≫ *τ*_1_, the inner sum is dominated by its largest value *z* = *N*_*m*_, leading to

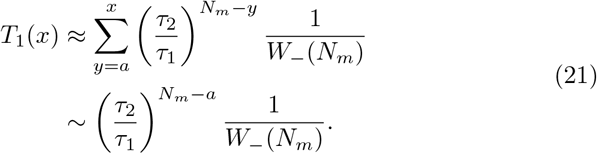

Given 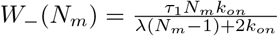, we obtain

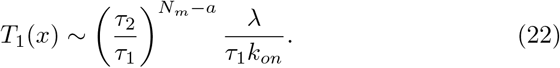

Figure 8(a) compares this approximation to the exact result from Eq. 10. The relative error remains below a factor of two already for *τ*_2_ ≳ 1.9 *τ*_1_ and is nearly independent of *N*_*m*_, *a*, and *b*.

**Fig 8.**
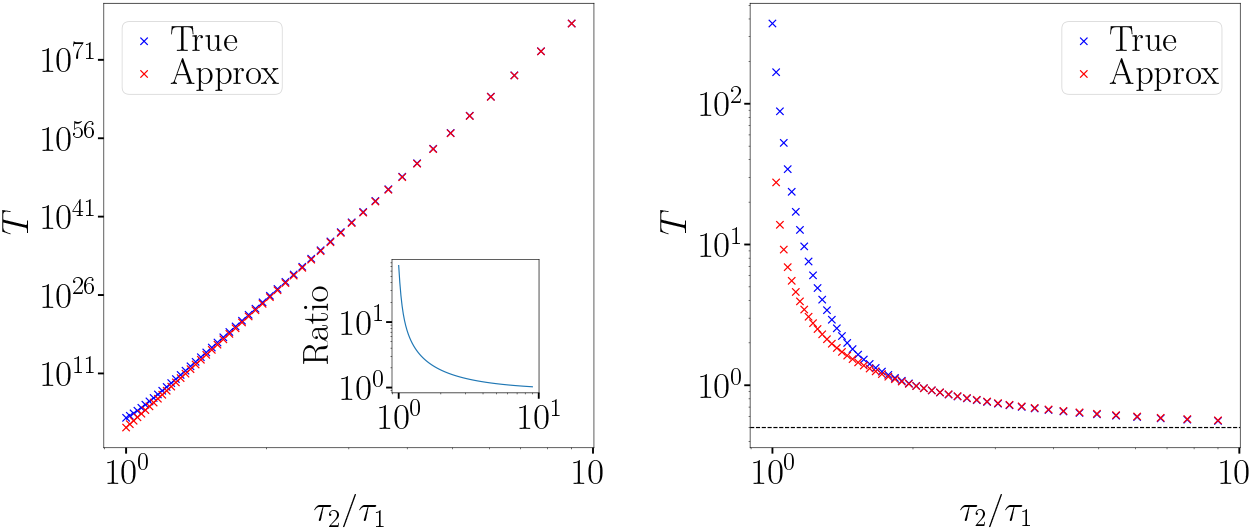
(a) Mean absorbing time of region 1 in the RR model. Exact result from Eq. 10 (blue) compared with the approximation Eq. 22 (red). Inset: ratio between the two. (b) Same for region 2. Approximation (red) from Eq. 25; dotted line: asymptotic form Eq. 24. Parameters: *N*_*m*_ = 100, *τ*_2_ = 9, *λ* = 0.01, *k*_*on*_ = 2.5 × 10^−4^, *a* = 10, *x* = 90.

– **In Region 2**.

Exchanging indices 1 and 2, we write

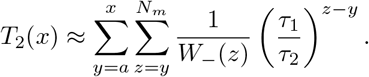

For *τ*_2_ ≫ *τ*_1_, the dominant contribution now comes from *z* = *y*, yielding

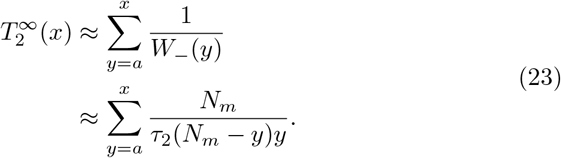

Evaluating the sum gives

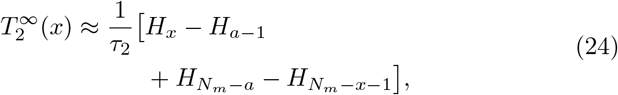

where 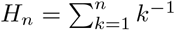 is the harmonic number.

To capture the finite *τ*_1_*/τ*_2_ corrections, we get,

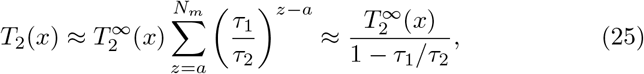

valid for *N*_*m*_ − *a* ≫ 1. Good agreement with the exact result is observed already for *τ*_2_ ≳ 2 *τ*_1_, see Fig. 8(b). Using the asymptotic expansion of the harmonic numbers,

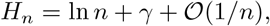

where *γ* ≃ 0.577 is the Euler–Mascheroni constant, one finds that 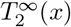 grows logarithmically. Combined with the geometric correction factor (1 − *τ*_1_*/τ*_2_)^−1^, this directly leads to the approximation in Eq. 12.

- To determine the growth of *T* (*x*) with *N*_*m*_ when *τ*_2_ is not necessarily much larger than *τ*_1_, the previously simple asymptotic approximation obtained for *τ*_2_ ≫ *τ*_1_ can no longer be used. Instead, we derive a general lower bound that holds for any *τ*_2_ *> τ*_1_.

For *N*_*m*_ ≥ *z > y >* 0,

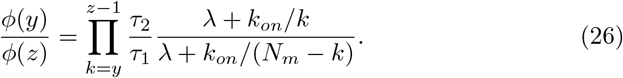

Taking logarithms gives

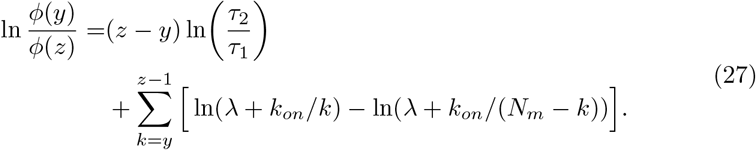

Define

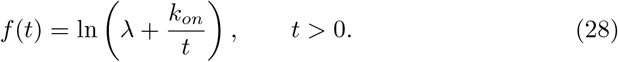

Since the function *f* is Lipschitz on [1, ∞), for all *k* ≥ 1,

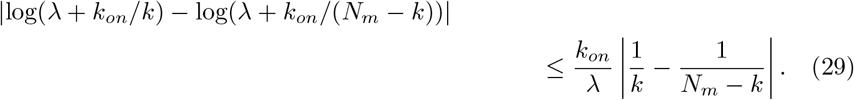

Summing from *k* = *y* to *z* − 1 yields

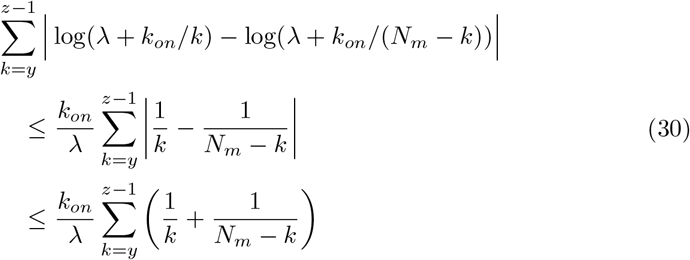

Using the bounds ln *n* ≤ *H*_*n*_ ≤ ln *n* + 1 for the *n*-th harmonic number *H*_*n*_, we obtain

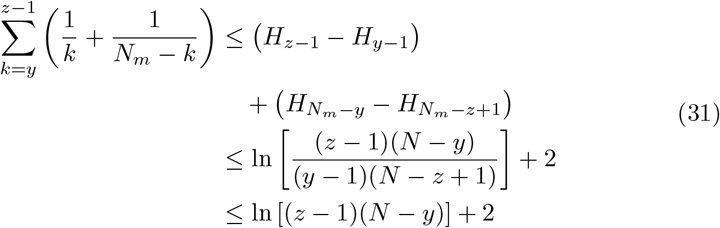

Combining this with (30) gives

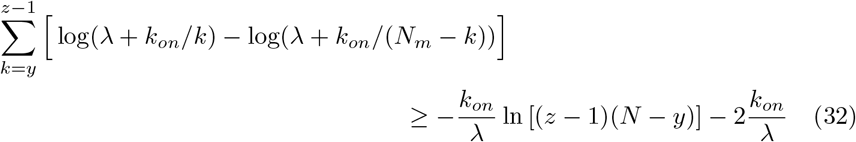

Substituting into (27), we obtain the lower bound

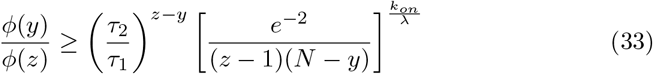

We now insert this estimate into Eq. 10:

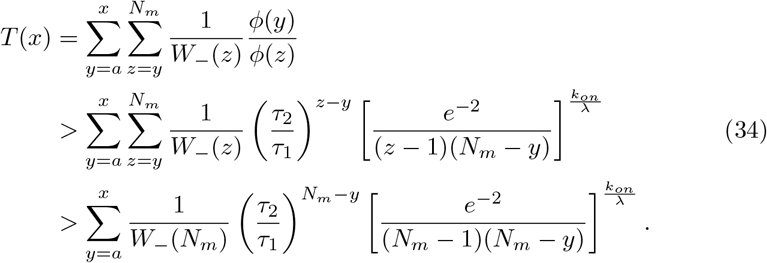

Approximating *W*_−_(*N*_*m*_) ≃ *τ*_1_*k*_*on*_*/λ* and using (*N*_*m*_ − *y*) ≤ (*N*_*m*_ − 1), we obtain

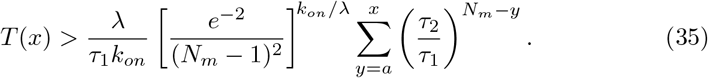

Evaluating the geometric sum yields

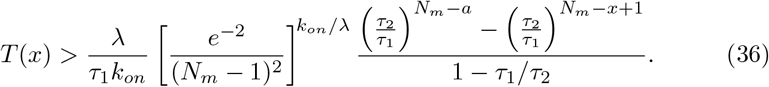

This expression shows that, for any *τ*_2_ *> τ*_1_, the dominant contribution to *T* (*x*) grows proportionally to 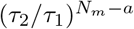, up to a polynomial prefactor in *N*_*m*_. Therefore, the mean absorbing time increases at least exponentially with *N*_*m*_ whenever *τ*_2_ *> τ*_1_.

- The case *τ*_1_ = *τ*_2_ requires a more careful treatment. A naive approximation,

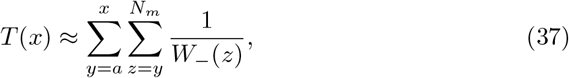

leads to

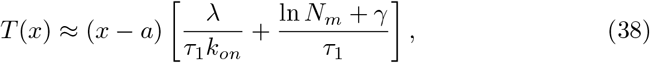

where *γ* is the Euler–Mascheroni constant. This predicts a logarithmic dependence on *N*_*m*_ that is not observed numerically, indicating that the approximation *ϕ*(*y*)*/ϕ*(*z*) ≃ 1 is insufficient in this regime.

We therefore return to the exact ratio

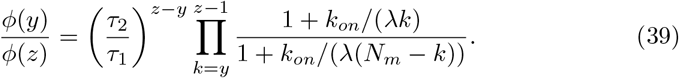

For *τ*_1_ = *τ*_2_, the prefactor is unity and the nontrivial contribution arises entirely from the product. Writing it in exponential form and expanding the logarithm to first order in *k*_*on*_*/λ*, we obtain

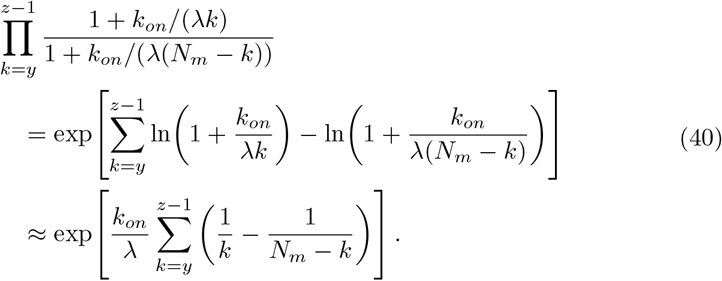

Evaluating the sums yields

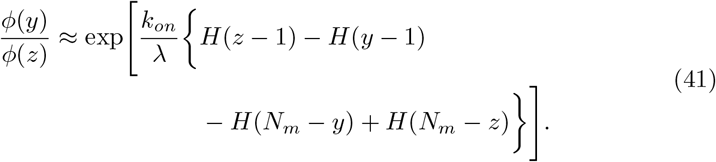

Using the asymptotic expansion *H*_*n*_ = ln *n* + *γ* + 𝒪(1*/n*), we obtain, for *z < N*_*m*_,

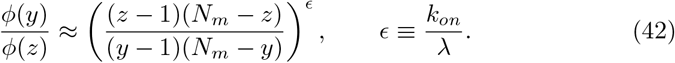

For the boundary term *z* = *N*_*m*_, the contribution *H*(*N*_*m*_ − *z*) must be set to zero, resulting in

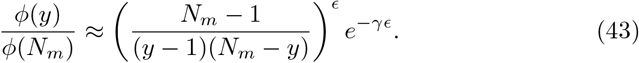

Substituting Eqs. (42) and (43) into Eq. 10, we obtain

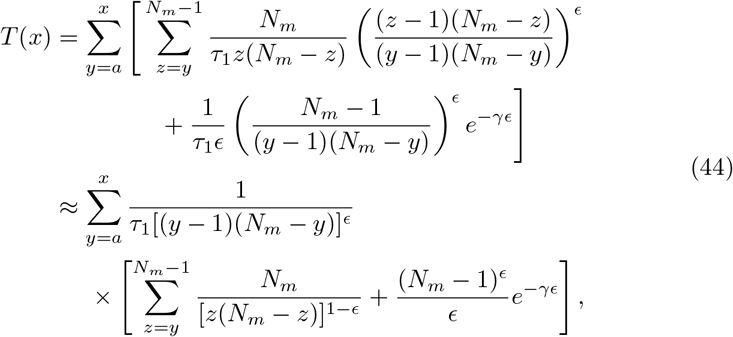

where we used (*z* − 1)^*ϵ*^*/z* ≃ *z*^*ϵ*−1^ as *z* will be always large, see later. The internal sum is evaluated using the Euler–Maclaurin formula,

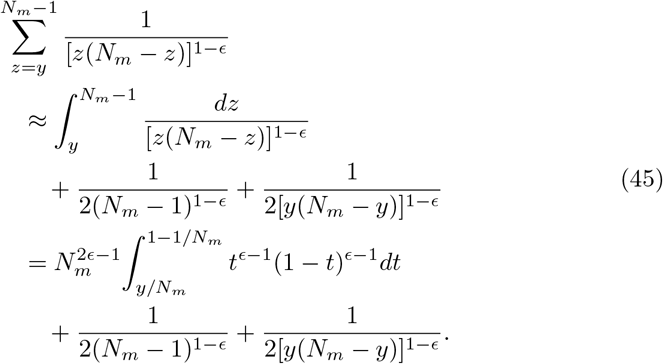

The integral is expressed in terms of the incomplete beta function 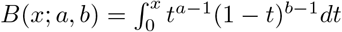,

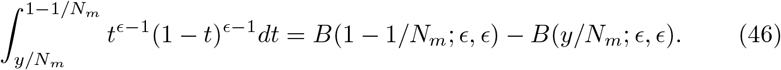

Using *B*(*ϵ, ϵ*) ∼ 2*/ϵ* as *ϵ* → 0 and

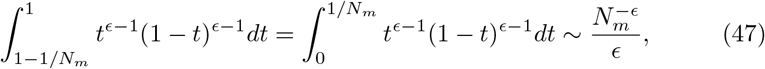

we obtain the bounds

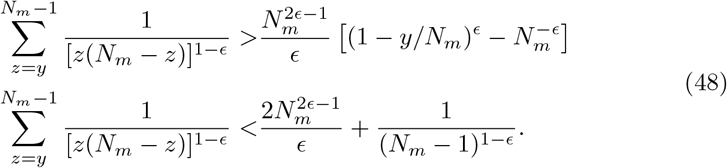

For *N*_*m*_ ≫ 1, this leads to the bounds

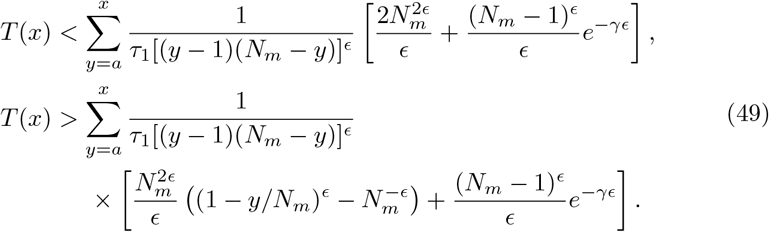

The remaining sum over *y* is estimated using the midpoint rule,

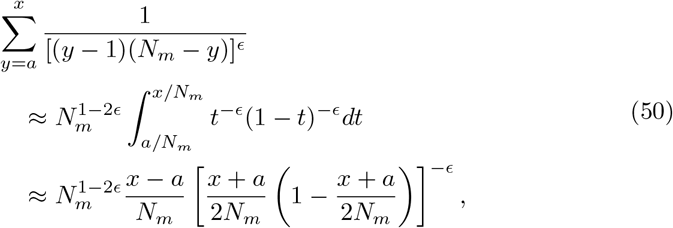

which is satisfactory provided *a/N*_*m*_, *x/N*_*m*_ is not too close from either 0 or 1. Here we typically use *a/N*_*m*_, *x/N*_*m*_ ∈ [0.1, 0.9]. Writing *a* = *α*_*a*_*N*_*m*_ and *x* = *α*_*x*_*N*_*m*_ with *α*_*a*_ + *α*_*x*_ = 1 to simplify the midpoint result, we finally obtain

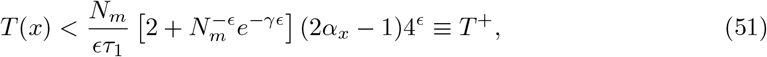

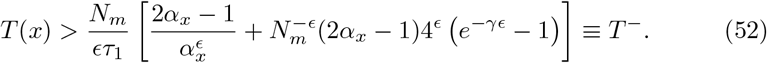

Importantly, equations (51) and (52) imply that the mean surviving time *T* grows linearly asymptotically with the total number of particles *N*_*m*_ when *τ*_1_ = *τ*_2_. This linear scaling contrasts with the logarithmic dependence predicted by the naive approximation discussed above.

In Fig. 9, we plot the exact mean surviving time *T* together with the upper and lower bounds *T* ^+^ and *T*^−^ as functions of *N*_*m*_. As expected from the looseness of the upper estimate in Eq. 51, the bound *T* ^+^ overestimates the true value of *T*. By contrast, the lower bound *T*^−^ provides a good approximation of *T* for large *N*_*m*_ for the parameters used.

**Fig 9.**
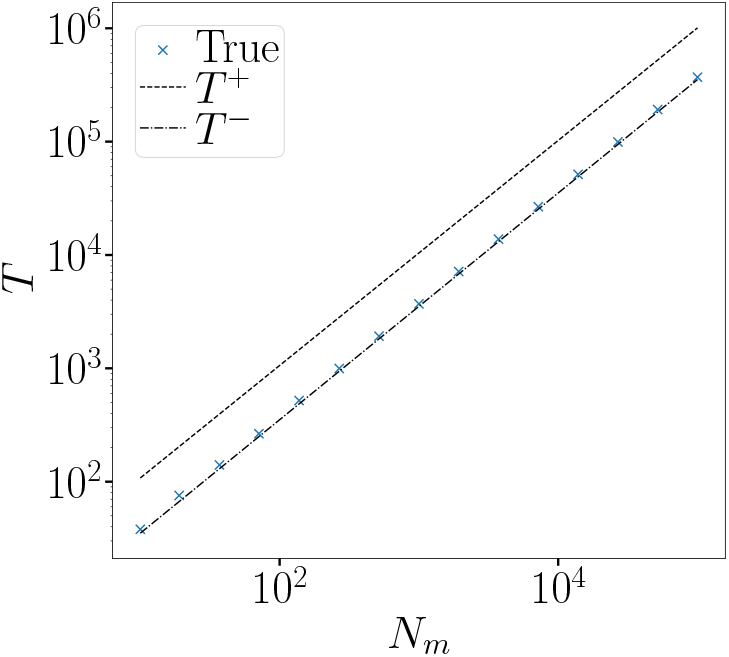
Mean surviving time *T* starting from *x* = 0.9*N*_*m*_ with an absorbing boundary at *a* = 0.1*N*_*m*_ for *τ*_1_ = *τ*_2_ = 9. The upper and lower bounds *T* ^+^ and *T*^−^ are given by equations (51) and (52). Other parameters are *λ* = 0.01 and *k*_*on*_ = 0.00025.

### S2 DEP

The diffusive epidemic process (DEP) is a discrete model as illustrated in Fig.1. We model the cell as a single chain, keeping only the dynamics on the membrane. Particles that should be diffusing inside the cytoplasm, called *c*, also diffuse along the chain at rate *D*_*c*_. Particles bind to the membrane, called *m*, diffuse at rate *D*_*m*_. We use *D*_*c*_ = 100*D*_*m*_ to reflect the fast diffusion in the cytoplasm. The coefficient 100 is coming from the ratio of the diffusion of the G protein in the cytoplasm and at the membrane [113]. However, the results do not depend qualitatively to a precise value of *D*_*c*_ as long as *D*_*c*_ *> D*_*m*_.

To make the correspondence between the continuous NDP, we discretize the system into *L* = 108 cells. Using more cells will not change the main findings. Each cell has a size Δ*x* = 27*/*108 = 0.25*µ*m. If the diffusion along the membrane in the NDP is *D* = 0.6 then, *D*_*m*_ = 0.6*/*Δ*x*^2^ = 9.6. The spontaneous unbinding rate *τ* and spontaneous binding rate *k*_*on*_ are identical. To obtain the recruitment rate that would lead to the same dynamics in the limit *D*_*c*_ → ∞, we take *λ*^*DEP*^ = *λ*^*NDP*^ *L*. Indeed, the rate at which a new particle is recruited in the DEP is *λ*^*DEP*^ *ρ*_*c*_ with *ρ*_*c*_ = *N*_*c*_*/L* the density of particle *c*. Then, the same recruitment rate in the DEP and NDP requires *λ*^*NDP*^ *N*_*c*_ = *λ*^*DEP*^ *N*_*c*_*/L* leading to *λ*^*DEP*^ = *λ*^*NDP*^ *L*.

### S3 Numerical simulations

Periodic boundary conditions are used unless stated otherwise.

#### Diffusive epidemic process (DEP)

Simulations of the diffusive epidemic process (DEP) [62, 63] are performed using the Gillespie algorithm [114, 115]. To incorporate spatial discretization, we employ the next-reaction method [116], which allows each lattice site to be associated with its own set of reaction channels. Spatial heterogeneities are thus implemented straightforwardly through site-dependent reaction rates.

#### Neutral drift polarity (NDP) model

For the neutral drift polarity (NDP) model [57, 59], reaction events are generated using the Gillespie algorithm [114, 115]. Between two reaction events separated by a time interval Δ*t*, all particles undergo independent Brownian displacements drawn from a Gaussian distribution with zero mean and variance 2*D*Δ*t*.

To reduce computational cost, particle positions are not updated after every individual reaction event. Instead, displacements are accumulated and applied only when the cumulative elapsed time satisfies ∑_*i*_ Δ*t*_*i*_ *> t*_th_. The threshold time *t*_th_ is chosen such that the typical diffusive displacement remains small compared to the characteristic spatial scales of the heterogeneities,

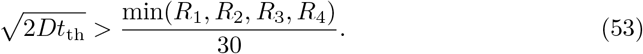

Because reaction rates depend on spatial position, this approximation renders the algorithm no longer strictly exact: during the interval [0, *t*_th_], a particle may temporarily be assigned to a region with incorrect local rates. We have verified that varying *t*_th_ within reasonable bounds does not qualitatively affect the results reported in this work.

#### Region–Cytoplasm–Region_*k*−1_ model

Simulations of the RCR model are performed using the Gillespie algorithm [114, 115], with reaction rates defined as in the main text.

### S4 Parameters

Unless stated otherwise, the parameters used for the NDP, RCR, and RR models are *λ* = 0.01 min^−1^, *τ* = 9 min^−1^, *k*_on_ = 5 × 10^−4^ min^−1^, and *D* = 1.2 µm^2^ min^−1^, following Ref. [59]. The rate *k*_on_ corresponds to the spontaneous binding of a cytoplasmic particle to a random membrane position. When two regions of equal size are considered, the corresponding rates are *k*_on,1_ = *k*_on,2_ = *k*_on_*/*2. The total number of particles *N* is varied and typically taken around 10^3^.

In Section Effect of disorder on two regions, the total one-dimensional system size is set to *L* = 50 µm.

In Section Emergence of oscillations, we choose *L*_1_ = *L*_3_ = 3.5 µm and *L*_2_ = *L*_4_ = 10 µm, consistent with the geometry of *Schizosaccharomyces pombe* cells [54, 87]. The spontaneous binding rates scale with region length, giving *k*_on,1_ = *k*_on,3_ ≈ 6.48 × 10^−5^ min^−1^ and *k*_on,2_ = *k*_on,4_ ≈ 1.85 × 10^−4^ min^−1^. The pole regions (R1 and R3) are slightly favored with *τ*_1_ = *τ*_3_ = 8.5 min^−1^, while *τ*_2_ = *τ*_4_ = 9 min^−1^.

In Sections Coexistence and Cell growth, unless stated otherwise, the DEP simulations use *L* = 108 with regions *L*_*R*1_ = *L*_*R*3_ = 14 and *L*_*R*2_ = *L*_*R*4_ = 40, *λ* = 1.08min^−1^, *τ*_1_ = *τ*_3_ = 8.5min^−1^, *τ*_2_ = *τ*_4_ = 9min^−1^, *D*_*c*_ = 960min^−1^, *D*_*m*_ = 9.6min^−1^, and *k*_on_ = 5 × 10^−4^min^−1^. The density *ρ* = *N/L* is varied, typically around *ρ* ≈ 15. These parameters correspond to the NDP model in the limit *D*_*c*_ → ∞.

In Section Cell growth, the system grows at a constant speed of including 2 sites per 20min up to doubling the initial system size, e.g., *L*_max_ = 216. Each added site contains either ⌊*ρ*⌋ or ⌊*ρ*⌋ + 1 to maintain the prescribed mean density *ρ*.

